# Sequence-Dependent DNA Base Selection Fidelity: A Kinetics-Based Model and its Validation

**DOI:** 10.64898/2026.07.30.741821

**Authors:** Koushik Ghosh, Parthasarathi Sahu, Shashikanta Barik, Hemachander Subramanian

## Abstract

DNA replication achieves error rates as low as 10^−9^-10^−11^ per base pair through the combined contribution of base selection, exonuclease proofreading, and mismatch repair. Among those processes, base selection determines the initial level of accuracy and is strongly modulated by local sequence context. Existing models address this dependence either by fitting individual rate constants for each sequence context or by invoking the global template properties, neither of which derives sequence dependence from the underlying thermodynamics and kinetics of base pair formation. Here we present a mechanism for sequence-dependent base selection fidelity, built from two physical properties: nearest-neighbor stacking thermodynamics and directional kinetic asymmetry. The model fits the experimentally observed mutation spectra from three mismatch repair-deficient organisms well (r=0.74, 0.70, and 0.58), and predicts that base-selection accuracy varies non-monotonically with temperature in a sequence-dependent manner. Our model, therefore, provides a framework that connects sequence-dependent thermodynamic and kinetic effects during nucleotide incorporation to experimentally observed mutation rates.

**Statement of significance:** The accuracy with which correct nucleotides are selected during DNA replication is dependent on the local sequence context, yet it remains poorly understood. Here we present a theoretical model in which the intrinsic base-selection error ratios emerge from the combined effect of nearest-neighbor base-stacking thermodynamics and a directional asymmetry in cooperativity between adjacent base pairs on the growing strand. We found that sequence-dependent error ratios calculated by the model exhibit correlations of r=0.74, 0.70, and 0.58 with mutation rates measured in mismatch-repair-deficient strains of three organisms. The model also predicts a non-monotonic dependence of base-selection accuracy on temperature. This work provides a theoretical framework, grounded in base-stacking energetics and asymmetric kinetics, for interpreting sequence-dependent base-selection error rates.

## A. Introduction

DNA replication occurs with extraordinary accuracy, with error ratios across organisms ranging from 10^*−*9^ to 10^*−*11^ [1, 2]. Such high fidelity arises from the coordinated action of three mechanisms [3–5]: (i) accurate selection of complementary nucleotides during the 5^*′*^ *→* 3^*′*^ polymerization step (base selection); (ii) selective removal of non-complementary nucleotides through the 3^*′*^*→* 5^*′*^ exonuclease proofreading activity; and (iii) post-replicative mismatch repair (MMR) of residual erroneous base-pairing. Each of these mechanisms contributes multiplicatively to overall accuracy [5, 6]: base selection alone reduces the error rate to *∼* 10^*−*4^ - 10^*−*5^ per nucleotide, exonuclease proofreading improves this by an additional 10 - 100-fold, and MMR suppresses the remaining error by 10^3^ *−* 10^4^ fold [3].

Because base selection is the first checkpoint for replication fidelity, it determines the intrinsic accuracy of nucleotide incorporation [5–7]. However, the accuracy of the base selection step is not uniform but varies with local DNA sequence context surrounding the nucleotide incorporation site [1, 5, 8–11]. This sequence dependence has also been demonstrated using high-throughput assays based on next-generation sequencing [12–15] and the Magnification via Nucleotide Imbalance Fidelity assay [16]. Despite demonstrating sequence-dependent fidelity, these studies provide little mechanistic understanding of how local sequence context influences nucleotide selection. Several theoretical models have been proposed to describe sequence-dependent base selection fidelity.

An early theoretical model to incorporate sequence dependence was proposed by Viljoen *et al*. [17]. In their model, the nucleotide insertion rate was modified according to the local GC content of the four preceding base pairs – an approach that reproduced the slower replication in GC-rich regions. P. Gaspard, in a series of works [18–20], further incorporated all 16 nearest-neighbor base-pair combinations into kinetic models and proposed an iterative algorithm, based on a backward inhomogeneous Markov chain approximation, to compute fidelity for a given template sequence. Building on this framework, Li *et al*. derived a fidelity expression (their Eq. 11) showing that nearest-neighbor template sequences dominate fidelity [21]. However, these approaches do not explicitly relate sequence-dependent fidelity to the thermodynamic stability and kinetic properties of local base-pair formation and dissociation.

Here, we propose an alternative approach in which sequence-dependent fidelity emerges from two physical properties: nearest-neighbor stacking thermodynamics [22] and asymmetric cooperativity [23–25]. Nearest-neighbor stacking determines the local thermodynamic stability of each base pair, whereas asymmetric cooperativity enables correctly formed base pairs to modulate the kinetic barriers at adjacent sites directionally.

Together, these mechanisms generate sequence-dependent nucleotide incorporation. In our previous work [26], we considered *sequence-independent* asymmetric cooperativity [23] and showed that, when combined with thermodynamic discrimination and irreversible covalent bond formation [27], it yields an error rate of *∼* 10^*−*4^ during daughter strand construction. The present study extends this by incorporating nearest-neighbor stacking thermodynamics and *sequence-dependent* asymmetric cooperativity into a continuous-time Markov chain [28, 29], and computes the error ratios for different sequence contexts.

We then correlate the calculated error ratios with the mutation accumulation data from MMR^*−*^ strains of *Bacillus subtilis, Mesoplasma florum*, and *Escherichia coli* [30]. In these strains, inactivation of mutL or mutS causes the loss of MMR activity, leading to mutations that show replication errors that escape both nucleotide selection and exonuclease proofreading [31]. Although proofreading reduces mutation rates [6, 8], the underlying sequence-dependent mutational biases are largely preserved [3, 32]. Thus, triplet-specific mutation rates may reflect underlying biases in nucleotide selection. Our study therefore provides a theoretical framework that connects sequence-dependent thermodynamics and kinetics of base-pair formation/dissociation with experimentally observed mutation patterns.

## B. Model Overview

To establish the theoretical foundation of our present extension, we first summarize the core framework of our previous model in [26]. In that model, error correction during daughter strand construction was described on a template consisting of *N* nucleotides. The central premise of that model is *sequence-independent asymmetric cooperativity*, a kinetic asymmetry in which a correctly formed base pair modifies the kinetic barriers for base-pair formation/dissociation at adjacent left and right neighboring sites in a directionally asymmetric manner. Specifically, a correct base pair at template position *m* (Fig. 1) lowers the kinetic barrier for basepair formation and dissociation at position *m* + 1, thereby catalyzing daughter-strand extension in the 3^*′*^ direction, while simultaneously raising the kinetic barrier for base-pair formation and dissociation at position *m −* 1, stabilizing the base pairs in the 5^*′*^ direction.

**Figure 1.**
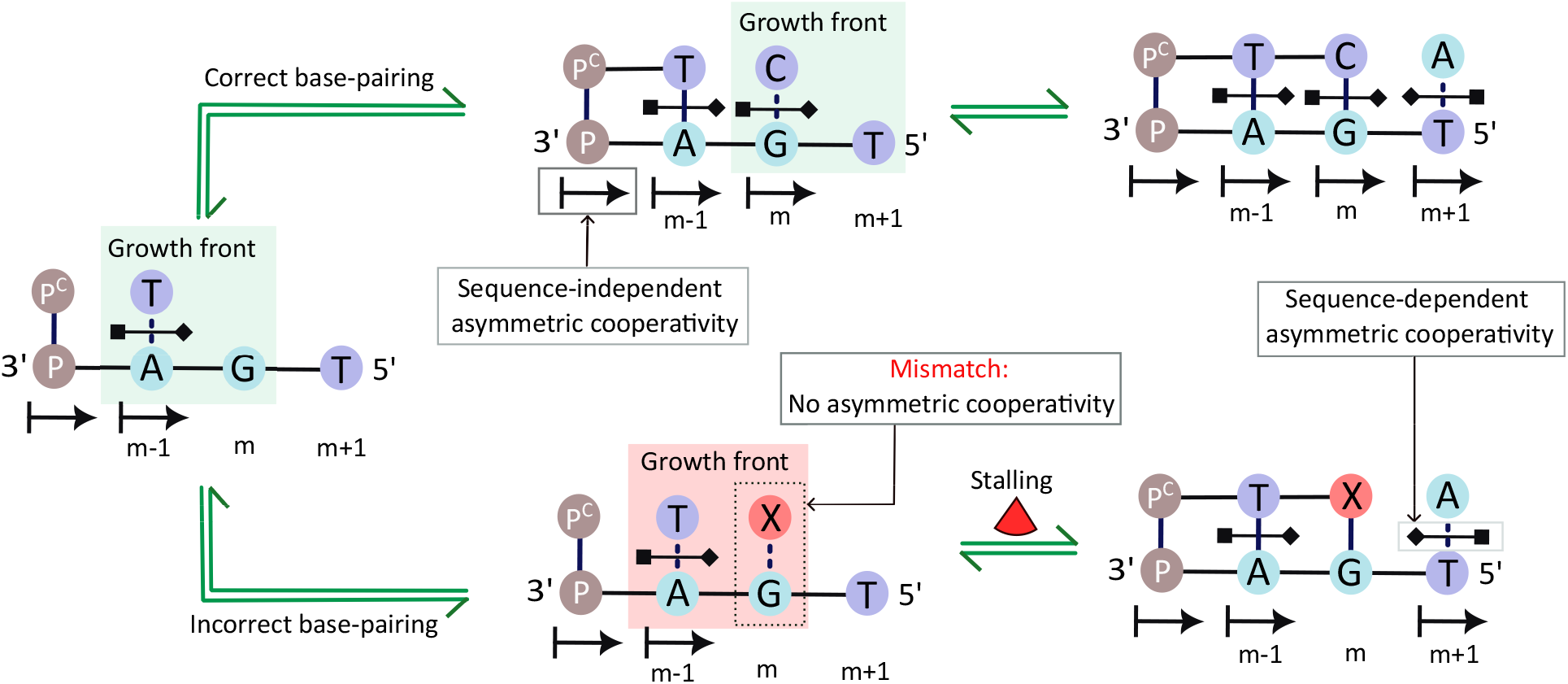
Schematic illustration of daughter-strand growth mediated by asymmetric cooperativity on a template strand of length *N* = 4. The upper pathway depicts correct base pairing, whereas the lower pathway depicts incorrect base pairing. A correctly formed base pair at position *m* directionally modulates the kinetic barriers for the formation and dissociation of neighboring base pairs through both sequence-independent and sequence-dependent asymmetric cooperativities. Sequence-independent asymmetric cooperativity, represented by the arrowhead and barhead symbols, lowers the kinetic barrier at position *m* + 1 and raises the barrier at position *m −* 1, thereby favoring unidirectional growth of the daughter strand. Sequence-dependent asymmetric cooperativity, represented by the square-diamond connector across the base pair, introduces an additional, smaller directional modulation of these neighboring kinetic barriers that depends on the identity of the base pair at position *m*. In contrast, a mismatch *X* at position *m* does not exert asymmetric cooperativity, resulting in the stalling of daughter-strand extension and destabilization of the preceding base pair, which initiates sequential unzipping.

In contrast, we assumed incorrect base pairs did not exhibit asymmetric cooperativity. An incorrect base pair at position *m* neither lowers the barrier for 3^*′*^ extension nor raises the barrier for 5^*′*^ stabilization. This has two consequences: (i) The absence of catalysis towards the 3^*′*^-end means subsequent nucleotide incorporation slows down. (ii) Simultaneously, the preceding base pair at *m −* 1 loses the stabilizing influence from its 3^*′*^-neighbor, leaving it prone to dissociate, which can initiate sequential unzipping of the previously formed base pairs. Consequently, the presence of an incorrect base pair introduces a time delay in daughter-strand construction compared to when there is no error, thereby giving rise to kinetic discrimination (Fig. 1).

In addition to this kinetic discrimination, correct and incorrect base pairs also differ in their intrinsic thermodynamic stability, resulting in different dissociation rates. In particular, incorrect base pairs dissociate faster than correct ones, with experimentally reported dissociation rates typically being about two orders of magnitude larger than those of correct base pairs [33]. Together, these kinetic and thermodynamic discriminations create a time lag between the correctly and incorrectly formed base pairs during reversible base-pairing stages of daughter strand construction. The contribution of this time lag to error correction depends on the covalent bond formation rate, which we assume is an irreversible step due to its highly exergonic nature (*∼* 12 *k*_*B*_*T*) [27, 34]. When the covalent bond formation rate is low, the reversible basepairing dynamics approach quasi-equilibrium [35] before covalent incorporation occurs. As a result, kinetic discrimination arising from the asymmetric cooperativity gets lost, and error correction is determined by the thermodynamic discrimination. In contrast, when covalent bond formation occurs on a time scale comparable to the base-pairing dynamics, covalent incorporation occurs before equilibration can be established, allowing both thermodynamic and kinetic discrimination to contribute to error correction [26].

While this framework demonstrates that error correction can emerge from the interplay of thermodynamic bias, kinetic asymmetry, and covalent-bond catalysis, it assumes a uniform asymmetric cooperativity factor and base-pair formation/dissociation rates that are independent of the template sequence. Consequently, the model predicts uniform error rates across different template sequences. Here, we extend this framework by incorporating sequence-dependent thermodynamics arising from nearest-neighbor base-pair stacking interactions [22] together with sequence-dependent kinetics associated with asymmetric cooperativity [24]. The resulting model introduces additional parameters describing the sequence-dependent stability and kinetic effects.

## C. Methods

We consider a template strand of *N* = 4 base pairs, where the base pair at the first position on the template 3^*′*^-end is always present and acts as a primer [36]. This base pair is assumed to be dynamically inactive – it neither dissociates nor participates in discrimination – and serves only to define the growth direction of the daughter strand. The remaining three sites constitute the active region over which base pair formation, dissociation, and error discrimination occur during the construction of the daughter strand.

To model the dynamics of strand construction, we use the continuous-time Markov chain [28, 29], in which each state represents a distinct base-pairing configuration along the template. Each site is represented by a variable taking values in *{* 0, 1, 2 *}* , corresponding to: (0) no base pair present, (1) a correct (complementary) base pair, or (2) an incorrect (mismatch) base pair. The full state space comprises all configurations consistent with these assignments. Transitions between states occur via base-pair formation and dissociation, with transition rates governed by local thermodynamic and kinetic parameters.

### I. Base-pair formation/dissociation and thermodynamic discrimination

Studies on short DNA duplexes report association rate constants *k*_*on*_ on the order of *≈* 10^6^ M^*−*1^sec^*−*1^ [37–39]. In our model, we consider the bare base pair formation at an unpaired site to occur with a sequence-independent rate *p* = 1 sec^*−*1^, given a nucleotide concentration of *≈* 1 *µM* [40, 41]. We assume that this bare base-pair formation rate is identical for correct and incorrect nucleotides; therefore, discrimination is entirely governed by dissociation. To incorporate sequence dependence into these dissociation rates, we evaluate the free energy of reversible base pairing using the nearest-neighbor thermodynamic formalism for DNA helix initiation and propagation at 1 M NaCl and 37^°^C [22]. Within this formalism, the stability of a base pair depends on both its own identity and its neighboring base pairs through stacking interactions. The corresponding nearest-neighbor thermodynamic parameters are listed in Table 1.

**Table 1:** Nearest-neighbor stacking and initiation parameters for DNA helix formation at 1 M NaCl, from [22]. 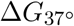 values are computed from Δ*G*^*°*^ = Δ*H*^*°*^ *− T* Δ*S*^*°*^ at *T* = 310.15 K (37^*°*^C).

| Nearest-neighbor pair | $\Delta H^\circ$ (kcal/mol) | $\Delta S^\circ$ (cal/mol·K) | $\Delta G_{37}^\circ$ (kcal/mol) |
| --- | --- | --- | --- |
| 5'-AA/TT-3' | -8.4 | -23.6 | -1.02 |
| 5'-AT/TA-3' | -6.5 | -18.8 | -0.73 |
| 5'-TA/AT-3' | -6.3 | -18.5 | -0.60 |
| 5'-CA/GT-3' | -7.4 | -19.3 | -1.38 |
| 5'-GT/CA-3' | -8.6 | -23.0 | -1.43 |
| 5'-CT/GA-3' | -6.1 | -16.1 | -1.16 |
| 5'-GA/CT-3' | -7.7 | -20.3 | -1.46 |
| 5'-CG/GC-3' | -10.1 | -25.5 | -2.09 |
| 5'-GC/CG-3' | -11.1 | -28.4 | -2.28 |
| 5'-GG/CC-3' | -6.7 | -15.6 | -1.77 |
| Initiation at G·C | 0 | -5.9 | +1.82 |
| Initiation at A·T | 0 | -9.0 | +2.8 |

When a base pair is isolated, dissociation is governed by initiation free energies Δ*G*_init_, which depend on the identity of the template nucleotide. Once at least one neighboring base pair is present, the duplex is already nucleated, and further stabilization arises solely from nearest-neighbor stacking interactions. In this stacked regime, initiation contributions do not enter the local stability, and dissociation is controlled exclusively by the local stacking free energy Δ*G*_stack_. The effective dissociation free energy is therefore defined as:

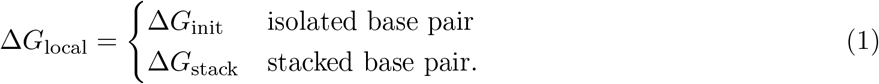

The dissociation rate of a correct base pair is then given by 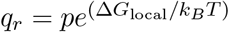. Incorrect base pairs are thermodynamically less stable and therefore dissociate more rapidly. We account for this by introducing a factor *f* = 10^2^ [42], such that the dissociation rate for an incorrect base pair is *q*_*w*_ = *f × q*_*r*_.

### II. Asymmetric cooperativity and kinetic discrimination

Nearest-neighbor thermodynamics governs the stability of individual base-pair configurations, but daughter-strand construction also requires both easy base-pair formation at the growing site and stabilization of previously formed base pairs [23]. To account for these competing requirements, we incorporate asymmetric cooperativity [23, 24]: a correctly formed base pair modulates the kinetic barriers of neighboring base pairs in a directionally asymmetric manner. Experimental studies have reported such intrinsic directional asymmetry in the unzipping of DNA duplex, including base pairs opening through asymmetric flipping pathways [43], different unzipping kinetics when identical duplexes are pulled from the 3^*′*^-or 5^*′*^-end [44], base-base coupling stronger in the 5^*′*^ *→*3^*′*^ than in the 3^*′*^ *→*5^*′*^ direction [45], and nanopore measurements revealing strand-orientation-dependent unzipping and translocation kinetics [46, 47].

In our previous work [26], we introduced the sequence-independent component of asymmetric cooperativity, wherein a pre-existing correct base pair lowers the kinetic barrier height for base-pair formation or dissociation at its neighboring site toward the template 5^*′*^-end (by a factor *α* = 5 *×* 10^3^), while raising the barrier height toward the 3^*′*^-end (by a factor *β* = 1*/α*). The chosen value of *α* is consistent with helix formation studies [37, 48, 49], which show that once the first base pair nucleates, each subsequent pair forms about 10^3^-fold faster. Likewise, kinetic analyses of measured polymerase rates have inferred barrier-height differences of *δ≈* 8 (T7) and *δ ≈*11 (pol*γ*) between correct and incorrect incorporation [50], corresponding to rate factors of 10^3^*−* 10^5^. These observations indicate that kinetic barriers of the magnitude implied by *α* are physically plausible in DNA copying systems.

Here, we additionally incorporate the sequence-dependent component of asymmetric cooperativity, whereby the modulation of these kinetic barriers depends on the orientation of the base pair [24, 25]. A correct base pair at the template position *m* exerts unequal effects on its two neighbors: the kinetic barrier at position *m* − 1 is modified differently from that at position *m* + 1, with the specific direction of this asymmetry determined by whether the base pair is 3^*′*^ *− G −* 5^*′*^*/*5^*′*^ *− C −* 3^*′*^ or 3^*′*^ *− C −* 5^*′*^*/*5^*′*^*− G −* 3^*′*^ (and analogously for A-T) (Fig. 2.A). Rotation of the base pair by 180^*°*^ reverses the direction of asymmetry. Incorrect base pairs, as in the sequence-independent case, are assumed not to exhibit sequence-dependent asymmetric cooperativity.

**Figure 2.**
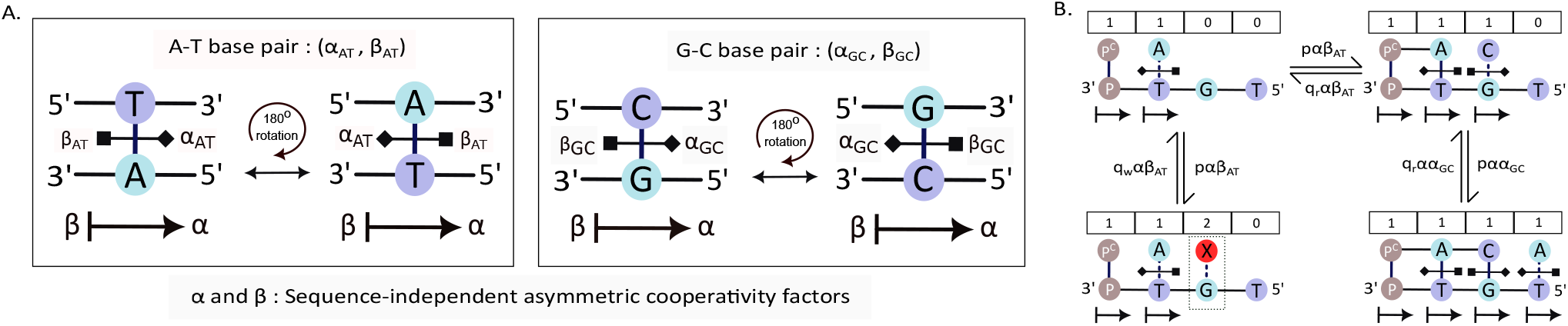
(A) Orientation dependence of sequence-dependent asymmetric cooperativity. A correctly formed base pair 3^*′*^ *− A −* 5^*′*^*/*5^*′*^ *− T −* 3^*′*^ modulates the kinetic barriers for the formation/dissociation of neighboring base pairs through the cooperativity factor (*α*_*AT*_ , *β*_*AT*_). Upon a 180^*°*^ rotation, the corresponding base pair 3^*′*^ *− T −* 5^*′*^*/*5^*′*^ *− A −* 3^*′*^ exhibits the same magnitude of cooperative effect, but with the direction of modulation reversed. An analogous orientation-dependent relationship holds for 3^*′*^ *− G −* 5^*′*^*/*5^*′*^ *− C −* 3^*′*^ and 3^*′*^ *− C −* 5^*′*^*/*5^*′*^ *− G −* 3^*′*^ base pairs, which are described by the cooperativity factors (*α*_*GC*_, *β*_*GC*_). (B) Illustrative state-transition diagram showing how the assigned sequence-independent and sequence-dependent asymmetric cooperativity parameters modify the transition rates of the model. States are represented by four-digit strings, where 0, 1, and 2 denote an unpaired site, a correctly paired base pair, and an incorrectly paired base pair, respectively. The intrinsic formation and dissociation rates *p, q*_*r*_, and *q*_*w*_ are multiplicatively modified by sequence-independent factors (*α, β*) and the sequence-dependent factors (*α*_*AT*_ , *β*_*AT*_ , *α*_*GC*_, *β*_*GC*_), thereby coupling local sequence context and strand orientation to the kinetics of daughter-strand growth.

We parameterize this effect through the base-specific cooperativity factors: for A-T pairs, left- and right-neighbor barriers are scaled by *α*_*AT*_ and *β*_*AT*_ ; for G-C pairs, by *α*_*GC*_ and *β*_*GC*_ (Fig.2A). These orientation-dependent modifiers act on the existing thermodynamic rates (*p,q*_*r*_, *q*_*w*_), which are already modified by the sequence-independent asymmetric cooperativity factors (*α, β*), providing sequence effects on the overall local extension kinetics during daughter strand construction (Fig. 2B).

### III. Covalent bond formation

As discussed above, covalent bond formation provides the irreversible step that preserves the kinetic discrimination generated during reversible base pairing. To incorporate this process into the model, we introduce the covalent bond formation rate *v*, which mediates the transition from fully base-paired configurations to the corresponding incorporated states. Since covalent bond formation is highly exergonic (*≈* 12*k*_*B*_*T*) [27, 34], the reverse transition is neglected.

Within the Markov chain, covalent bond formation is represented by absorbing transitions from fully base-paired states (1111/1121) to their respective irreversible product states (R/W). The transition rate from these states to these covalent states is parameterized by the dimensionless covalent bonding rate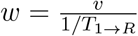, where *T* _1*→R*_ is the conditional mean-first passage time from the initial state 1000 to the fully paired correct state R. Since 1*/T*_1*→R*_ represents the characteristic rate of daughter-strand construction through reversible base-pairing dynamics, the dimensionless *w* quantifies the covalent bond formation rate relative to this rate.

### IV. Error rates

The dynamics of daughter-strand construction are determined by the transition rate matrix Q, whose off-diagonal elements represent allowed transitions between different base-pairing configurations, while the diagonal elements ensure probability conservation [28, 29]. The time evolution of the probability vector P(t), defined over all configurations, is governed by

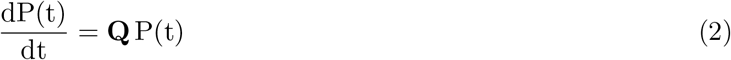

With the covalent bond formation described above, the system evolves toward absorbing states (R/W) corresponding to correct and incorrect incorporation. The steady-state probability vector P^SS^ is given by:

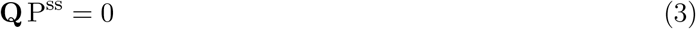

The fidelity of the daughter-strand construction is governed by the steady-state probabilities of absorption into the covalently stabilized correct and incorrect states, denoted by 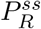 and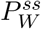, respectively. The intrinsic base-selection error ratio is therefore defined as

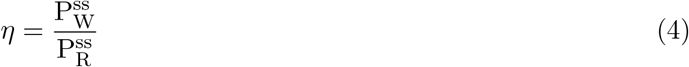

This ratio provides a measure of the intrinsic fidelity arising from nucleotide incorporation, independent of subsequent proofreading and mismatch repair.

### V. Experimental Verification

We computed the intrinsic base-selection error ratio (*η*) for each template strand 3^*′*^ *− PN*_*r*_*MN*_*l*_ *−* 5^*′*^, with *N*_*r*_, *M, N*_*l*_ = *A, G, T, C*. The base pair at the first template position acts solely as a primer and does not participate in discrimination. The incorrect base pair is located at position *m* = 3, while the neighboring positions *m−*1 and *m*+1 define the local sequence context. For a triplet sequence oriented as 3^*′*^*−N*_*r*_*MN*_*l*_ *−*5^*′*^, the correctly paired daughter strand is 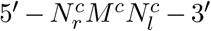. An incorrect incorporation at the central position corresponds to 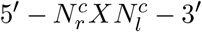, where *X ≠M*^*c*^. We then compute the intrinsic error ratio *η*, as defined in the Methods section, for each triplet context.

Thereafter, to validate the sequence-dependent trends predicted by the model against experimental observations, we consider mutation-accumulation studies in MMR^*−*^ organisms [30]. Since the MMR operates after nucleotide misinterpretation and removes replication errors, its absence exposes the underlying mutation spectrum generated during DNA replication [51]. The mutation rates reported in such strains (10^*−*8^ per triplet per generation) are several orders of magnitude lower than the intrinsic base-selection error ratios *η* (10^*−*4^ *−* 10^*−*5^), reflecting the action of polymerase proofreading, which improves replication fidelity by approximately 10^2^ *−* 10^3^ fold [6, 8]. Nevertheless, the underlying sequence-dependent biases are largely preserved [3, 32]. Consequently, the relative variation in mutation rates across triplet contexts in MMR^*−*^ strains is expected to correlate with the relative variation in base-selection error rates. We therefore evaluate proportional correspondence between model predictions and experimental observations via log-log correlation.

## D. Results

### I. Sequence-Dependent Base-Selection Fidelity

We first examined the effect of sequence context on the intrinsic base-selection error ratio *η* by incorporating nearest-neighbor base-stacking thermodynamics and *sequence-independent* asymmetric cooperativity into the model. Triplets were grouped into strong (*S ≡{G, C }*) and weak (*W≡ {A, T}*) classes based on the thermodynamic stability of their constituent base pairs [52]. To identify the value of the dimensionless covalent bond formation rate *w* at which both the kinetic and thermodynamic discrimination operate synergistically, we plot the dependence of the class-averaged error ratio *η* on *w* (Fig. 3A). For *w <* 1, corresponding to *v <* 1*/T*_1*→R*_, the system approaches quasi-equilibrium among the base-pairing states. Under this condition, the kinetic discrimination is lost, and error correction is governed solely by thermodynamic discrimination. As *w >* 1, i.e., *v >* 1*/T*_1*→R*_, covalent bond formation occurs before equilibration is established, allowing kinetic discrimination to contribute to error correction, and thereby reducing *η*. We therefore selected *w* = 10^4^ for the subsequent analysis, where both thermodynamic and kinetic discrimination contribute to error correction. This choice assumes that covalent bond formation is much faster than the strand construction speed. The corresponding class-averaged error ratios *η* at this value of *w* are shown in Fig. 3B.

**Figure 3.**
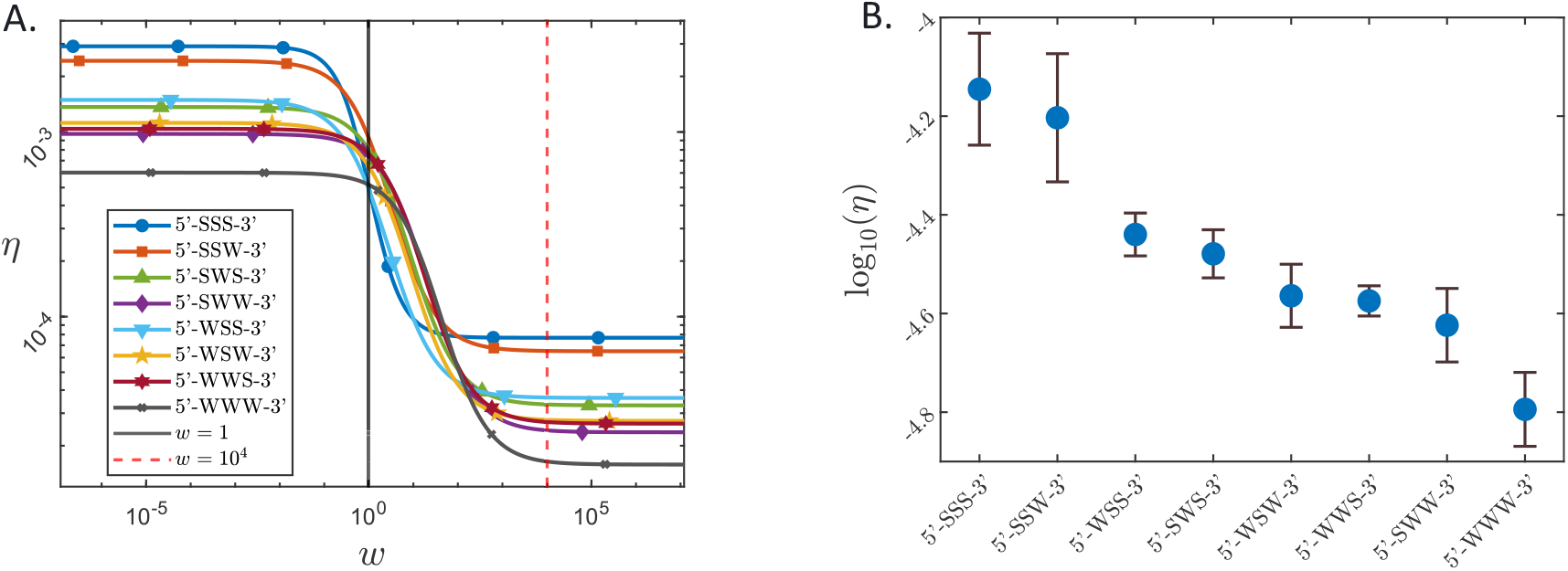
Base-selection error ratio *η* across nearest-neighbor triplet contexts classified by strong (*S ≡ {G, C}*) and weak (*W ≡{A, T}*) base composition. Each triplet label (e.g., 5^*′*^ *−SSS −*3^*′*^, 5^*′*^ *−WWW −*3^*′*^) denotes a distinct combination of flanking bases surrounding the central position where error base-pairing occurs. (A) Dependence of base selection error ratio *η* on the dimensionless covalent bond formation rate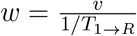. Curves represent mean error ratios for triplet classes grouped by strong (*S ≡ {G, C}*) and weak (*W ≡ {A, T }*) base composition. As *w* increases, the system transitions from a thermodynamic regime (*w <* 1), in which *η* is high, to a thermodynamic-plus-kinetic regime (*w >* 1), where kinetic discrimination becomes effective and *η* decreases. The vertical solid line indicates *w* = 1; the dashed line marks *w* = 10^4^, the value used in subsequent calculations. (B) Base-selection error ratios at *w* = 10^4^ for the corresponding triplet classes. Error bars denote standard deviations within each triplet group. The mean error ratio decreases systematically with increasing A/T content: the all-strong triplet class (5^*′*^ *−SSS −*3^*′*^) exhibits the highest *η*, the all-weak class (5^*′*^ *−WWW −*3^*′*^) the lowest, and the intermediate classes display progressively lower error ratios as the fraction of weak base pairs increases.

The plot in Fig. 3B shows a striking pattern: the strongest triplet group (5^*′*^ *−SSS −*3^*′*^) exhibits the highest intrinsic base selection error rate *η*, whereas the weakest group (5^*′*^ *−WWW −*3^*′*^) shows the lowest, with intermediate groups displaying *η* values that increase monotonically with G/C content. This variation suggests that base-stacking thermodynamics constitutes a primary determinant of intrinsic base-selection error rates *η* [42, 53]. In A-T-rich (5^*′*^ *−WWW −*3^*′*^) sequence contexts, new base pairs, whether correct or incorrect, are weakly bound to the template strand, allowing an incorrect base to dissociate more freely before the covalent bond is formed. Conversely, in G-C-rich (5^*′*^ *−SSS−* 3^*′*^) contexts, the stronger stacking interactions of each G-C pair increase the residence time of new base pairs, thereby increasing the probability that covalent bond formation occurs before dissociation, even when the pairing is incorrect. Therefore, the flanking base composition modulates the retention of incorrect base pairs through its effect on base-pair stability: A-T-rich contexts allow more frequent dissociation, whereas G-C-rich contexts tend to stabilize the incorrect base pair and increase the probability that the incorrect base pair is permanently incorporated during daughter strand construction.

This finding aligns with the analysis of *Petruska et al*. [54], who showed that “nucleotide misinsertion frequencies are determined by 5^*′*^ nearest-neighbor base stacking,” and that the local G.C content influences error correction efficiency. Their Fig.1 quantifies this effect, showing that the incorporation efficiency of a mismatched base analogue (2-aminopurine) increases monotonically with the proportion of G.C base pairs flanking the insertion site.

These results suggest that the nearest-neighbor stacking thermodynamics is an important determinant of the sequence-dependent variation in model-predicted base selection error ratios. However, the biological relevance of this variation depends on whether the predicted differences in base selection fidelity are reflected in experimentally observed mutation patterns. If nearest-neighbor thermodynamic stacking is a crucial aspect of the sequence dependence formed during the base selection step [8, 55, 56], then triplet contexts predicted by the model to exhibit larger intrinsic base selection error ratios should also display higher mutation frequencies experimentally.

Therefore, we next compared the model-predicted base selection error ratios *η* with context-dependent mutation rates *µ* measured in mutation-accumulation (MA) experiments of mismatch repair-deficient (MMR^*−*^) strains of *Mesoplasma florum, Bacillus subtilis*, and *Escherichia coli* [30]. Although experimentally observed mutation rates (*∼*10^*−*8^) are several orders of magnitude lower than the predicted base selection error ratios (*∼*10^*−*4^ *−* 10^*−*5^) due to proofreading, the underlying sequence-dependent pattern is largely preserved [3, 32]. Fig. 4 shows a log-log plot of the model-predicted intrinsic base selection error ratio (*η*) against the experimentally observed mutation rates (*µ*) across triplet contexts.

**Figure 4.**
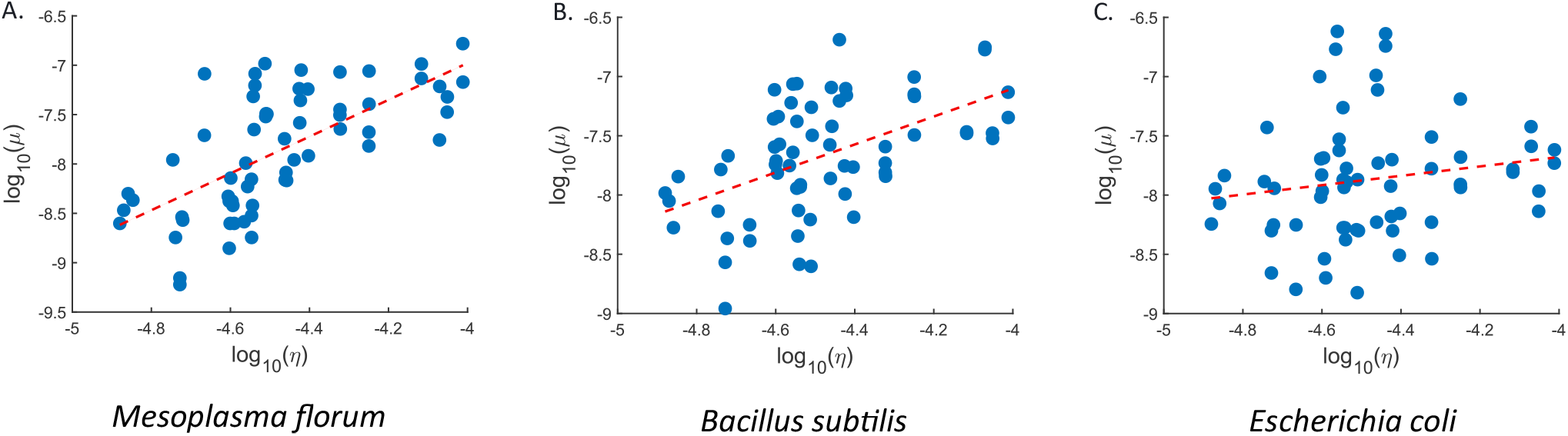
Log-log comparison of model-predicted base-selection error ratios *η* with mutation rates *µ* measured in mutation accumulation (MA) experiments of MMR^*−*^ strains. Each panel shows log_10_(*µ*) versus log_10_(*η*) for (A) *Mesoplasma florum*, (B) *Bacillus subtilis*, and (C) *Escherichia coli*. The model incorporates nearest-neighbor stacking thermodynamics and sequence-independent asymmetric cooperativity, without orientation-dependent effects. The corresponding Pearson correlation coefficients are 0.67, 0.52, and 0.17 for *Mesoplasma florum, Bacillus subtilis*, and *Escherichia coli*, respectively. The correlation is strongest for *Mesoplasma florum* and decreases for *Bacillus subtilis* and *Escherichia coli*.

Across all three organisms, the log-log plots in Fig. 4 show positive correlations between the model-predicted base selection error ratios and the experimentally observed mutation rates. The Pearson correlation coefficients are 0.67 for *Mesoplasma florum*, 0.52 for *Bacillus subtilis*, and 0.17 for *Escherichia coli*. These positive correlations suggest that nearest-neighbor thermodynamic stacking, together with the sequence-independent asymmetric cooperativity, captures an important component of sequence-dependent variation in mutation rates. The strongest correlation is obtained for *Mesoplasma florum*, whereas weaker correlations are obtained for *Bacillus subtilis* and *Escherichia coli*. This decrease in correlation suggests that thermodynamic stacking alone cannot fully account for the observed mutation rates. In the current model, triplets with the same stacking composition are assigned identical kinetic properties, regardless of orientation. As a result, the model does not incorporate sequence-dependent effects arising from strand orientation and their influence on local kinetic barriers. We therefore next incorporated *sequence-dependent* asymmetric cooperativity [24] into the model, allowing triplets with identical stacking compositions but different orientations to exhibit distinct kinetic properties. We then examined whether this improves the agreement between the predicted base selection error ratios *η* and the experimentally observed mutation rates *µ*.

As described in the methods section, the incorporation of sequence-dependent asymmetric cooperativity suggests that each correctly formed base pair exerts an orientation-dependent influence on neighboring sites, modulating the kinetic barriers for base-pair formation and dissociation. The corresponding kinetic parameters, Θ = *{α*_*AT*_ , *β*_*AT*_ , *α*_*GC*_, *β*_*GC*_*}* , quantify the orientation-specific cooperative effect associated with AT and GC base pairs (Fig. 2A). We treat these parameters as free parameters and optimized for each organism individually using a genetic algorithm that minimizes the weighted root-mean-square error (RMSE) [57] between the model-predicted base selection error rates *η* and experimentally observed mutation rates *µ*. The results obtained after incorporating sequence-dependent asymmetric cooperativity with optimized parameters are shown in Fig. 5.

**Figure 5.**
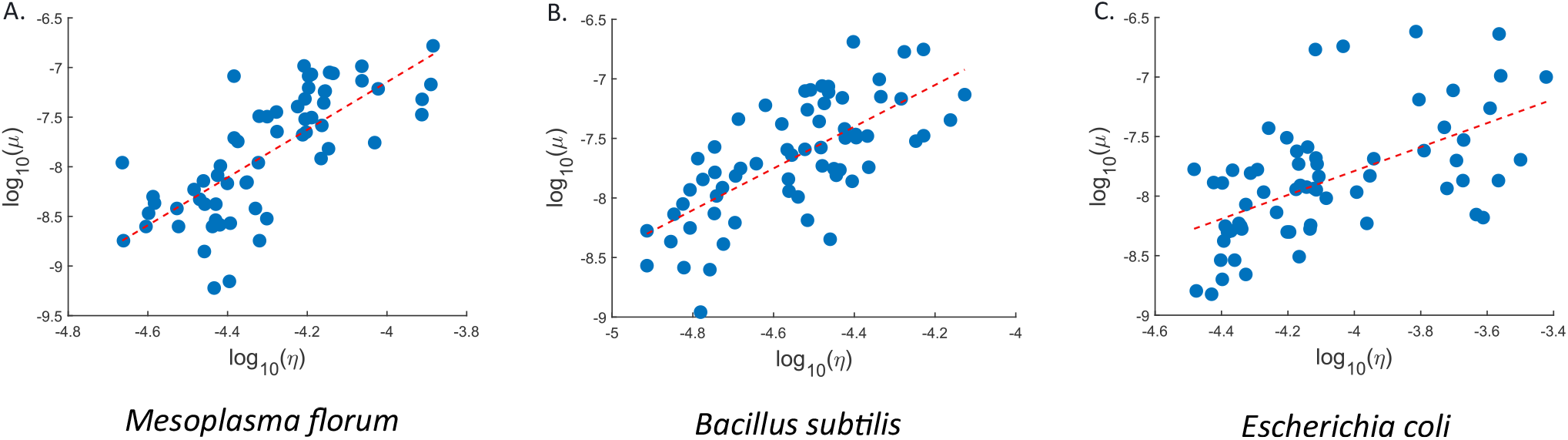
Log-log comparison of model-predicted base-selection error ratios *η* with mutation rates *µ* measured in mutation accumulation (MA) experiments of (MMR^*−*^) strains, after incorporating sequence-dependent asymmetric cooperativity. Each panel shows log_10_(*µ*) versus log_10_(*η*) for (A)*Mesoplasma florum*, (B) *Bacillus subtilis*, and (C) *Escherichia coli*. The orientation-specific sequence-dependent asymmetric cooperativity parameters (Θ = *{α*_*AT*_ , *β*_*AT*_ , *α*_*GC*_, *β*_*GC*_*}*), were optimized independently for each organism. The resulting Pearson correlation coefficients are 0.74, 0.70, and 0.58, respectively.

Table 2 shows the Pearson correlation before and after incorporating sequence-dependent cooperativity (SDAC). In all three organisms, incorporating sequence-dependent asymmetric cooperativity increases the correlations between the model’s predicted base selection error ratios *η* and the experimental mutation rate *µ*, suggesting that orientation-dependent kinetics may influence context-dependent base selection errors. The strongest correlation is observed for *Mesoplasma florum* (*r* = 0.74) followed by *Bacillus subtilis* (*r* = 0.70) and *Escherichia coli* (*r* = 0.58). Notably, *Mesoplasma florum* is a minimal organism completely lacking MMR machinery [58]; as a result, its mutation spectrum most accurately reflects intrinsic polymerase behavior. On the other hand, the *Bacillus subtilis* and *Escherichia coli* strains still possess some MMR activity, as they are deficient in only one component (MutS or MutL) [30]. This partial MMR activity may obscure the true effect of base-selection accuracy on mutation patterns.

**Table 2:** Correlation between model-predicted base-selection error ratios (*η*) and experimentally measured mutation rates (*µ*) before and after incorporating sequence-dependent asymmetric cooperativity (SDAC).

| Organism | Without SDAC |  |  | With SDAC |  |  |
| --- | --- | --- | --- | --- | --- | --- |
| | $r$ | $R^2$ | $p\text{-value}$ | $r$ | $R^2$ | $p\text{-value}$ |
| *Mesoplasma florum* | 0.67 | 0.44 | $3.29 \times 10^{-09}$ | 0.74 | 0.55 | $5.14 \times 10^{-12}$ |
| *Bacillus subtilis* | 0.52 | 0.27 | $1.01 \times 10^{-05}$ | 0.70 | 0.48 | $1.75 \times 10^{-10}$ |
| *Escherichia coli* | 0.17 | 0.03 | $1.83 \times 10^{-01}$ | 0.58 | 0.34 | $5.15 \times 10^{-07}$ |

The optimized sequence-dependent asymmetric cooperativity parameters for each organism are shown in Table 3. In the model, nearest-neighbor stacking thermodynamics determine the local stability of the base-pairing configuration, while sequence-independent asymmetric cooperativity introduces a common directional kinetic bias. Together, they establish the primary differences among triplet contexts. Sequence-dependent asymmetric cooperativity introduces an additional orientation-specific kinetic contribution. This introduces a further source of sequence dependence beyond that arising from stacking thermodynamics and sequence-independent asymmetric cooperativity. Although the fitted parameter values differ among organisms, incorporating this effect improves the model’s ability to distinguish among triplet contexts.

**Table 3:** Optimized sequence-dependent asymmetric cooperativity parameters for each organism.

| Organism | $\alpha_{AT}$ | $\beta_{AT}$ | $\alpha_{GC}$ | $\beta_{GC}$ |
| --- | --- | --- | --- | --- |
| *Mesoplasma florum* | 0.41 | 0.45 | 0.87 | 0.71 |
| *Bacillus subtilis* | 4.05 | 2.92 | 2.28 | 1.78 |
| *Escherichia coli* | 13.26 | 10.97 | 5.04 | 20.05 |

### II. Temperature Dependence of Base-Selection Fidelity

Since the dissociation rate is given by 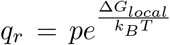, with Δ*G*_*local*_ = Δ*H*^*°*^ *− T* Δ*S*^*°*^ and Δ*H*^*°*^ and Δ*S*^*°*^ are assumed to be temperature-independent [59, 60], the dissociation rates *q*_*r*_ and *q*_*w*_ = *fq*_*r*_ vary explicitly with temperature. This implies that the intrinsic error ratio *η*, which depends upon the rates of correct and incorrect base-pair dissociation, is also temperature-dependent. Therefore, we next examine the temperature dependence of *η* across different triplet contexts. For illustration, we use the optimized parameter set of *Mesoplasma florum*, which exhibits the highest correlation with experimental mutation rates. Fig. 6 shows the variation of 1*/η* with temperature for each triplet class. All triplet classes exhibit a non-monotonic trend: 1*/η* initially increases with temperature, reaches a maximum at intermediate temperature, and then decreases at higher temperature. Furthermore, both the magnitude of 1*/η* and the temperature at which it occurs vary among triplet classes, indicating that local base-pair stability influences the temperature at which discrimination is maximized.

**Figure 6.**
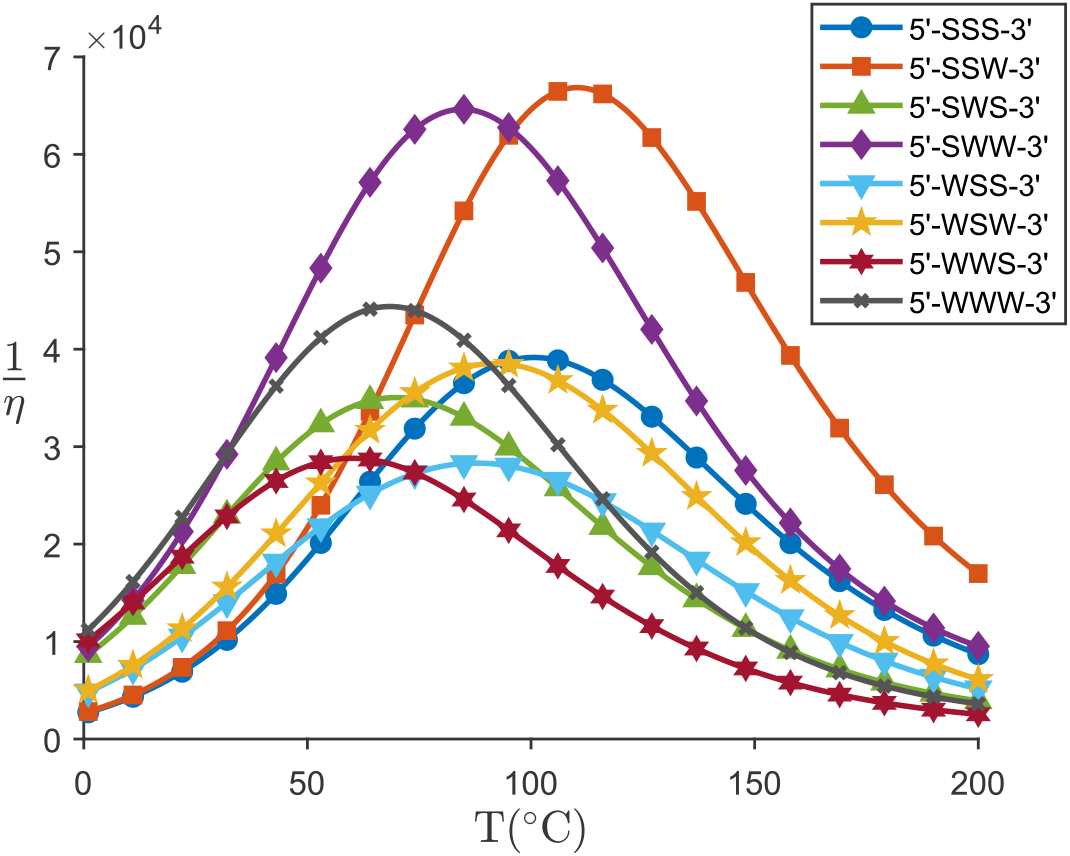
The plot shows the base-selection fidelity (1*/η*) as a function of temperature (*T*) for triplet groups classified by their strong (*S ≡ G, C*) and weak (*W* = *A, T*) base composition. The calculation uses the sequence-dependent cooperativity parameters optimized for *Mesoplasma florum*. Although the magnitude of fidelity varies among triplet groups, all groups show a common non-monotonic dependence on T: fidelity initially increases, reaches a maximum at an intermediate T, and decreases at higher T.

This variation arises from the competition between thermal destabilization of base pairs and the enhanced dissociation of incorrect base pairs relative to correctly paired bases. At low temperatures, both correct and incorrect base pairs are highly stable, suppressing dissociation and thereby limiting the opportunities for error correction. As temperature increases, the stability of both classes of base pairs decreases. However, incorrect base pairs dissociate 100-fold faster than correct base pairs; they dissociate preferentially, increasing the probability that incorrect base pairs are rejected before irreversible covalent bond formation occurs. Consequently, 1*/η* increases, reaching a maximum at an intermediate temperature where discrimination between the correct and incorrect base pair is most effective. At higher temperatures, even correctly paired bases become unstable, reducing the distinction between correct and incorrect incorporation. As a result, the efficiency of error correction decreases and 1*/η* declines.

The predicted non-monotonic temperature dependence aligns with the observation of Xue *et al*. [61], who reported that the substitution error rate of the KleLF and T4 polymerases exhibits non-monotonic variation with temperature. Furthermore, Das *et al*. [62] demonstrate that biological copying mechanisms inherently achieve maximal accuracy in an intermediate temperature range. Our results extend these observations by showing that the temperature of optimal accuracy varies with local sequence context, due to the thermodynamic stability of base pairs and a kinetic barrier modulated by asymmetric cooperativity. This suggests that temperature-dependent modulation of base-selection fidelity can serve as a physical mechanism for tuning mutation patterns, potentially enabling adaptive responses to changing environmental conditions [6].

## E. Discussion

DNA replication achieves error rates as low as 10^*−*9^ to 10^*−*11^ per base pair through a hierarchy of correction mechanisms [5, 6]. The first step of this hierarchy is base selection - the initial nucleotide incorporation step - which establishes the baseline accuracy and is strongly influenced by local sequence context [3, 11]. Our study addresses how this sequence context modulates base-selection fidelity by integrating thermodynamic and kinetic determinants of base pair formation and relating them to experimentally observed mutation patterns [30].

The central finding is that nearest-neighbor stacking thermodynamics [22], along with sequence-independent asymmetric cooperativity [23], governs the baseline magnitude of context-dependent error rates during base selection, while sequence-dependent asymmetric cooperativity [24, 25] modulates the kinetic barriers differently at the 5^*′*^ and 3^*′*^ neighboring sites depending on base pair orientation. This enables contexts such as 5^*′*^ *−* GC *−* 3^*′*^ and 5^*′*^ *−* CG *−* 3^*′*^ to exhibit distinct error rates despite identical thermodynamic stability. This separation of thermodynamic and kinetic contributions provides insight into a long-standing question about the accuracy of base selection: whether sequence-dependent variation arises from base pair stability in duplex [63, 64] or from polymerase interaction at the active site [65]. Our model shows that such variation can emerge from base-pair thermodynamics and directional kinetic coupling between adjacent base pairs during strand extension.

The positive correlation between the model-predicted intrinsic base selection error rates and mutation rates in MMR^*−*^ strains of three organisms [30] supports the roles of nearest-neighbor thermodynamics, sequence-independent and sequence-dependent asymmetric cooperativities in context-dependent base selection. The strongest correlation observed for *Mesoplasma florum* is consistent with its nature as a minimal organism without functional mismatch repair machinery [58]. Consequently, the mutation spectrum is likely to provide a more direct representation of DNA polymerase’s intrinsic fidelity. The comparatively weaker correlation in *Bacillus subtilis* and *Escherichia coli* likely reflects residual mismatch repair activity in singlegene knockouts (mutS or mutL) and their more complex replication machinery [30]. Two triplet contexts with zero observed mutations in the *Mesoplasma florum* dataset (supplemental information Table S1) were excluded from correlation analysis. These zeros likely reflect sampling limitations arising from finite mutation accumulation generations and the rarity of these triplets in the *Mesoplasma florum* genome [30].

Our approach to sequence dependence differs in how parameters are assigned. Rather than fitting separate rate constants for each sequence context [17, 18, 21], we derived context from nearest-neighbor stacking free energies - whose influence on fidelity is well established [54, 64, 66] - coupled with four organism-specific sequence-dependent asymmetric cooperativity parameters. These parameters are optimized to match the experimental mutation rates, but they correspond to a physical quantity: the asymmetric change in kinetic barrier height for base pair formation and dissociation at neighboring sites. This framework allows the model to capture diverse sequence contexts without individual calibration. Furthermore, base-selection fidelity exhibits a non-monotonic dependence on temperature (Fig. 6). To fully characterize this dependence, we explored a broad temperature range, as living organisms maintain genetic information fidelity across vastly different thermal conditions that would denature many macromolecules [67, 68]. Examining how the fidelity of base selection varies with temperature may therefore help elucidate the physical basis underlying this robustness.

However, in our model we treat all mismatches as energetically equivalent through a single destabilization factor *f* = 10^2^[42], although experimental studies show that different mismatches can have different discrimination factors. For example, G.G mismatches are more stable than T.T or A. A mismatches in the same sequence context [64]. Since complete thermodynamic parameters for internal mismatches are available [69, 70], incorporating mismatch-specific discrimination factors may improve the model. Also, we consider only the nearest-neighbor interactions and do not account for longer-range sequence effects that have been reported in [21, 71]. Extending the framework to include the long-range sequence dependence may provide a more complete description of sequence-dependent fidelity.

## Statements and Declarations

### Data availability

All datasets and code used in this study are publicly available at the GitHub repository: https://github.com/KoushikGPhysics/Sequence-dependent-base-selection.

## Author Contributions

H.S. conceived the project and obtained the grants. K.G. coded and analysed the results with the help of P.S., S.B., and H.S. All four authors contributed to the writing, reviewing, and editing of the manuscript.

## Acknowledgments

Support for this work was provided by the Science & Engineering Research Board (SERB), Department of Science and Technology (DST), India, through a Core Research Grant with file no. CRG/2020/003555 and a MATRICS grant with file no. MTR/2022/000086.

## Supplemental Information

### A Mutation Data from Mismatch Repair-Deficient Strains

Context-dependent base-substitution mutation rates per triplet per generation in both replichores of *Mesoplasma florum, Bacillus subtilis*, and *Escherichia coli* mismatch-repair-deficient (MMR^*−*^) mutation accumulation lines from [30] on a 10^*−*8^ scale.

**Table S1:** Context-dependent mutation rates per triplet per generation for *Mesoplasma florum*.

| 5'-end | Target Pre-mutation Site |  |  |  | 3'-end |
| --- | --- | --- | --- | --- | --- |
|  | T | C | A | G |  |
| T | 0.07 | 2.23 | 0.34 | 6.24 | T |
|  | 0.18 | 2.61 | 0.47 | 4.40 | C |
|  | 0.27 | 4.83 | 0.25 | 8.21 | A |
|  | 0.30 | 5.79 | 0.25 | 8.93 | G |
| C | 1.10 | 2.26 | 0.25 | 7.35 | T |
|  | 0.00 | 4.04 | 0.82 | 1.75 | C |
|  | 0.18 | 3.56 | 0.38 | 10.30 | A |
|  | 0.59 | 2.10 | 0.68 | 6.10 | G |
| A | 0.50 | 3.02 | 0.29 | 1.95 | T |
|  | 0.14 | 1.21 | 0.70 | 3.23 | C |
|  | 0.43 | 10.38 | 0.06 | 8.19 | A |
|  | 0.42 | 5.73 | 0.38 | 3.20 | G |
| G | 0.26 | 4.78 | 0.42 | 3.13 | T |
|  | 0.00 | 6.75 | 1.80 | 1.52 | C |
|  | 1.02 | 3.35 | 0.72 | 8.53 | A |
|  | 1.10 | 16.54 | 0.69 | 8.73 | G |

**Table S2:** Context-dependent mutation rates per triplet per generation for *Bacillus subtilis*.

| 5'-end | Target Pre-mutation Site |  |  |  | 3'-end |
| --- | --- | --- | --- | --- | --- |
|  | T | C | A | G |  |
| T | 0.27 | 0.26 | 0.89 | 1.17 | T |
|  | 4.18 | 1.02 | 4.39 | 7.91 | C |
|  | 2.14 | 0.74 | 1.04 | 1.22 | A |
|  | 8.71 | 1.76 | 1.81 | 6.93 | G |
| C | 0.73 | 1.44 | 2.68 | 3.41 | T |
|  | 2.29 | 3.21 | 8.08 | 16.86 | C |
|  | 1.64 | 2.56 | 4.59 | 3.31 | A |
|  | 8.63 | 7.08 | 3.80 | 17.66 | G |
| A | 0.53 | 0.25 | 0.43 | 0.56 | T |
|  | 2.54 | 0.65 | 1.14 | 5.50 | C |
|  | 1.43 | 0.62 | 0.11 | 0.41 | A |
|  | 7.73 | 1.72 | 0.45 | 3.19 | G |
| G | 1.77 | 3.00 | 1.53 | 1.86 | T |
|  | 6.22 | 4.51 | 2.65 | 9.90 | C |
|  | 6.00 | 3.33 | 1.94 | 1.55 | A |
|  | 20.46 | 7.37 | 1.38 | 6.80 | G |

**Table S3:** Context-dependent mutation rates per triplet per generation for *Escherichia coli*.

| **5'-end** | Target Pre-mutation Site |  |  |  | **3'-end** |
| --- | --- | --- | --- | --- | --- |
|  | T | C | A | G |  |
| T | 0.50 | 0.42 | 1.13 | 1.68 | T |
|  | 1.35 | 0.66 | 10.00 | 1.99 | C |
|  | 1.14 | 0.53 | 0.57 | 1.29 | A |
|  | 0.53 | 1.19 | 1.48 | 0.50 | G |
| C | 1.30 | 0.29 | 0.20 | 1.65 | T |
|  | 2.96 | 1.16 | 7.72 | 3.78 | C |
|  | 3.72 | 0.59 | 0.29 | 1.56 | A |
|  | 2.38 | 1.23 | 1.86 | 2.58 | G |
| A | 0.85 | 0.15 | 0.56 | 0.16 | T |
|  | 2.02 | 0.70 | 5.47 | 1.35 | C |
|  | 1.46 | 0.51 | 0.22 | 0.56 | A |
|  | 0.96 | 0.31 | 1.16 | 0.50 | G |
| G | 17.02 | 1.08 | 2.06 | 1.67 | T |
|  | 23.00 | 2.41 | 10.24 | 6.46 | C |
|  | 24.07 | 0.73 | 1.08 | 3.09 | A |
|  | 18.12 | 1.86 | 0.59 | 2.09 | G |

